# Breaking the Phalanx: Overcoming Bacterial Drug Resistance with Quorum Sensing Inhibitors that Enhance Therapeutic Activity of Antibiotics

**DOI:** 10.1101/2025.01.17.633658

**Authors:** Jon-Michael Beasley, Dorjbal Dorjsuren, Sankalp Jain, Marielle Rath, Ricardo Scheufen Tieghi, Alexander Tropsha, Anton Simeonov, Alexey V. Zakharov, Eugene Muratov

## Abstract

Antibiotic-resistant bacterial infections loom over humanity as an increasingly deadly threat. There exists a dire need for new treatments, especially those that synergize with our existing arsenal of antibiotic drugs, to help overcome the gap in antibiotic efficacy and attenuate the development of new antibiotic resistance in the most dangerous pathogens. Quorum-sensing systems in bacteria drive the formation of biofilms, increase surface motility, and enhance other virulence factors, making these systems attractive targets for the discovery of novel antibacterials. Quorum-sensing inhibitors (QSIs) are hypothesized to synergize with existing antibiotics, making bacteria more sensitive to the effects of these drugs. In this study, we aimed to find the synergistic combinations between the QSIs and known antibiotics to combat the two deadliest hospital infections - *Pseudomonas aeruginosa* and *Acinetobacter baumannii.* We mined biochemical activity databases and literature to identify known, high-efficacy QSIs against these bacteria. We used these data to develop and validate a Quantitative Structure-Activity Relationship (QSAR) model for predicting QSI activity and then employed this model to identify new potential QSIs from the Inxight database of approved and investigational drugs. We then tested binary mixtures of the identified QSIs with 11 existing antibiotics using a combinatorial matrix screening approach with ten (five of each) clinical isolates of *P. aeruginosa* and *A. baumannii*. Amongst explored drug combinations, 22 exhibited a synergistic effect. Although no mixture inhibiting all the strains was found, piperacillin combined with ketoprofen, indomethacin, and piroxicam demonstrated the broadest antimicrobial action. We anticipate that further preclinical investigation of these combinations of novel repurposed QSIs with a known antibiotic may lead to novel clinical candidates.

**Table of Content Graphic:** 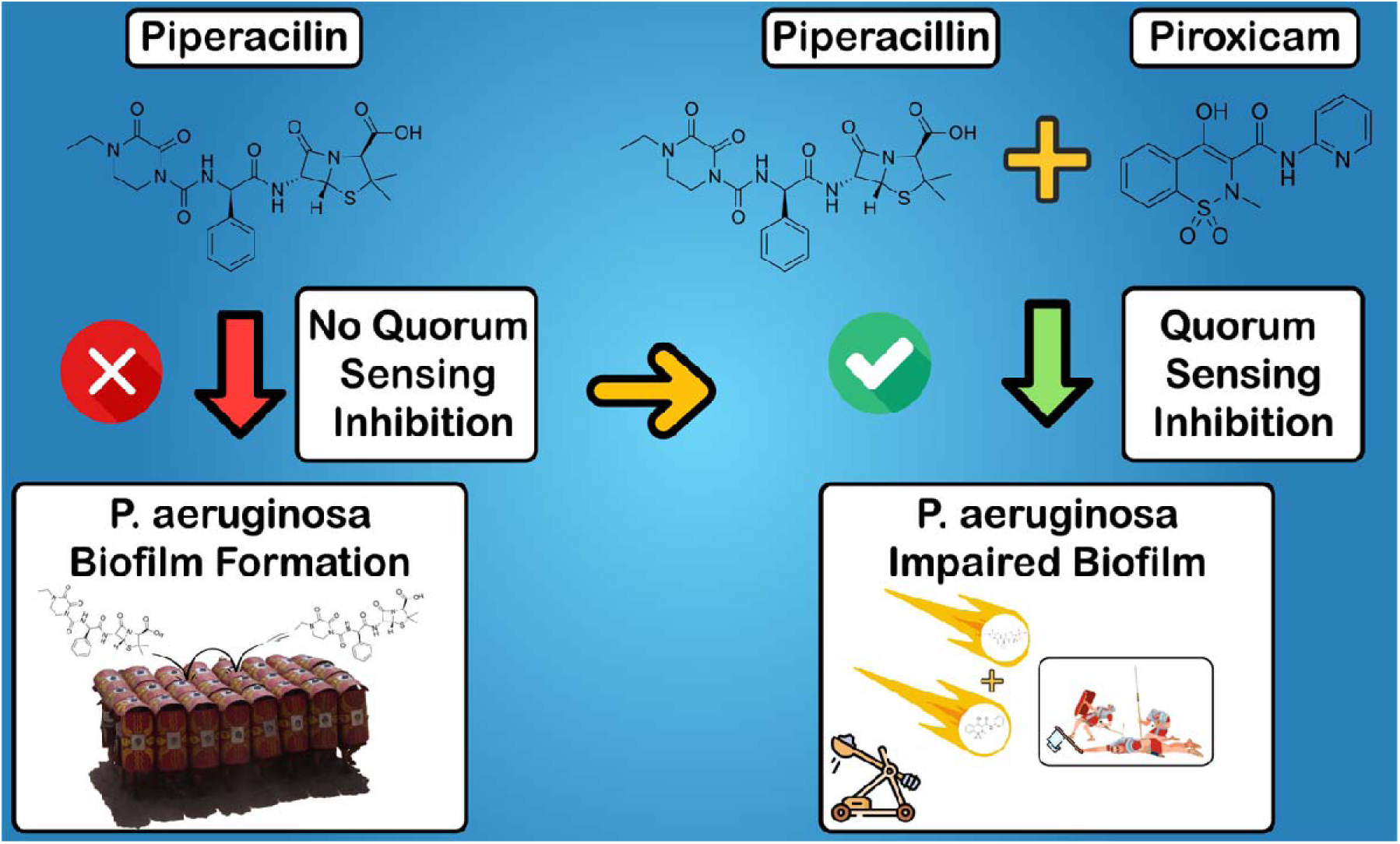

## Introduction

Antibiotic resistance, a global crisis of formidable proportions, has catapulted scientific communities into an urgent quest for innovative strategies to address the escalating threat to public health.^1^ Most current antibiotics have been designed to directly kill pathogenic bacteria by destroying cell membranes or interfering with protein synthesis, triggering “Life or Death selection pressure and promoting the evolution of microbial resistance. Indeed, almost all pathogenic bacteria are resistant to commonly used antibiotics, which effectively renders frontline treatments ineffective.^2^ The extensive use of antibiotics has led to the creation of “superbugs” that can resist a wide variety of common antibiotics.^3^ A report by the British Government estimated that by 2050, antimicrobial resistance could cause 10 million deaths each year (**Figure 1**), rivalling cancer as a global cause of death, and cause a cumulative loss of US $100 trillion to world GDP.^4^

**Figure 1.**
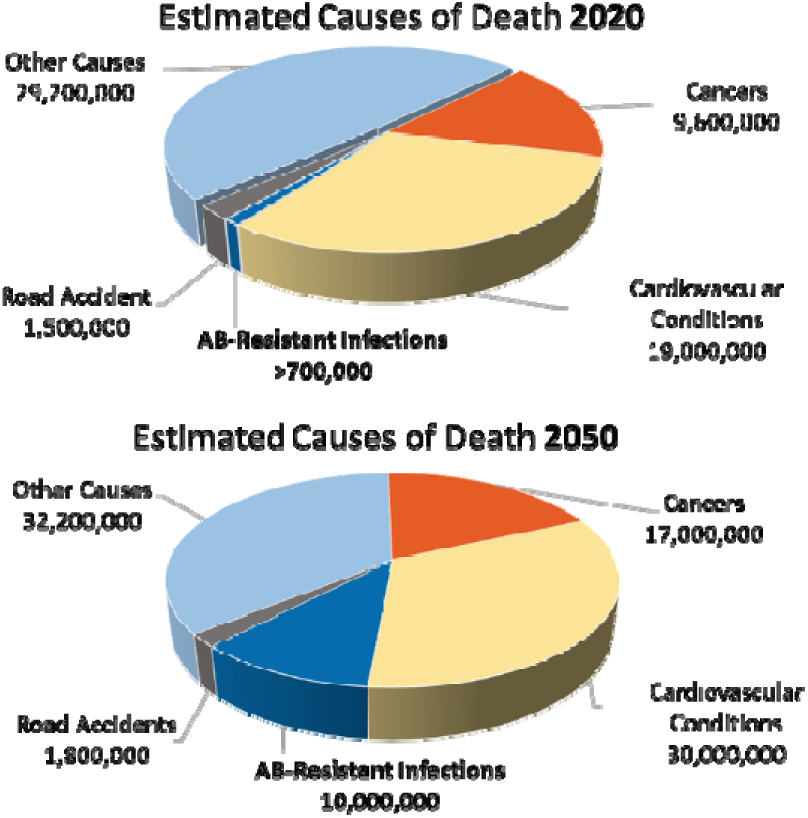
Antibacterial Resistance (ABRI) expected to become a major cause of death by 2050. ABRI-related deaths are expected to increase disproportionately, making up nearly 11% of total deaths in 2050 compared to an estimated 1% in 2020.^5,6^

As we face the antibiotic resistance crisis, where even routine infections pose a menacing risk, the development of novel solutions is imperative.^7^ Despite the urgent need for new treatment options, the speed of antibiotic development lags far behind the rate at which bacteria are evolving to become multidrug resistant (MDR). The lack of innovation on this front can be attributed to the high cost of $1 billion^8^ but low success rates of new drugs, as well as to the high likelihood that bacteria will eventually develop resistance to monotherapies. These expectations of low potential profitability effectively de-incentivize pharmaceutical companies to work on discovering and manufacturing such drugs.^9^

Fortunately, existing antibiotics can still be leveraged against bacterial infections. Combination therapy with synergistic antibiotics having distinct mechanisms of action and different targets shows promise in reducing antibiotic resistance. In addition, dual therapies can more effectively and quickly treat bacterial infections.^10^ Furthermore, lower doses of each antibiotic in the dual therapy can be used, which reduces the risk of toxicity and adverse effects.^11^

Several combination therapies are already on the market, such as amoxicillin/clavulanate (Augmentin), tazobactam/piperacillin (Zosyn), and trimethoprim/sulfamethoxazole (Bactrim). Other antibiotic combinations have shown success in experiments, such as vancomycin + trimethoprim and vancomycin + nitrofurantoin against *Escherichia coli*^12^, polymixin B + meropenem + ampicillin/sulbactam against *Acinetobacter baumannii*^13^, and polymixin B + aztreonam + amikacin against colistin-resistant *E. coli*^14^. Interestingly, several non-antibiotic compounds have synergistic effects with select antibiotics, including ursolic acid/oleanolic acid + ampicillin/oxacillin against MRSA^15^ and plant-derived flavonoids catechol-type flavonoid-7,8-dihydroxyflavone, myricetin, and luteolin + colistin against MDR bacteria^16^.

It is important to note that combination therapies do not necessarily need to be specially formulated, but rather each compound can be administered separately. Because combination therapies are less common, there are few dosing guidelines; therefore, physicians and pharmacists play a vital role when it comes to dosing in the clinic.^17^

### Quorum Sensing Inhibitors and their Synergy with Antibiotics

Among the diverse avenues of research aimed at circumventing antibiotic resistance, the inhibition of quorum sensing (QS) has emerged as a promising and relatively unexplored frontier.^4,18,19^ Unlike traditional antibiotics that directly target essential cellular processes, QS inhibitors (QSIs) seek not to eliminate bacteria outright but disarm them, diminishing their ability to mount a coordinated defense.^20^ At the heart of this exploration lies the recognition that as bacterial populations reach a critical density, they release signaling molecules into their environment.^20,21^ These molecules serve as communal messages, allowing bacteria to gauge their numbers and orchestrate collective actions, such as forming biofilms, enhancing surface motility, or activating virulence factors, which have been associated with increased resistance to common antibiotics.^20,21^ Disrupting this communication breaks their ability to act as a united front, potentially rendering them more susceptible to the host immune system and traditional antibiotics.^22,23^

The potential for synergistic interventions that simultaneously target both QS and traditional antibiotic pathways holds the promise of a two-pronged assault on bacterial resilience.^19^ Synergistic combinations involving QS inhibitors may lower the required doses of each drug and help avoid adverse events while exploiting the greater therapeutic effect.^24,25^ For example, low-dose gentamicin combined with amoxicillin improves treatment of bacterial endocarditis.^26,27^ Furthermore, synergistic combinations have the potential to rescue the efficacy of antibiotics for which bacteria have previously developed resistance, enabling the practice of “antibiotic recycling” of the first line-of-defense drugs.^28^

### Focusing on the Threat of Drug-Resistant *Pseudomonas aeruginosa* and *Acinetobacter baumannii* Infections

Multidrug resistant ESKAPE pathogens, such as *Enterococcus faecium, Staphylococcus aureus, Klebsiella pneumoniae, Acinetobacter baumannii, Pseudomonas aeruginosa and Enterobacter spp.* have been recognized as emerging threats to public health. Among these 6 bacteria, the World Health Organization (WHO) prioritizes *Acinetobacter baumannii, Pseudomonas aeruginosa* as critical for R&D of new antibiotics. Antibiotic resistance in *P. aeruginosa* poses a challenge due to the variety of resistance mechanisms, complicating the selection of effective treatment. An analysis of bloodstream infections from a major US hospital revealed higher mortality rates associated with *P. aeruginosa* isolates exhibiting a ‘difficult to treat’ resistance phenotype (DTR), which is characterized by resistance to fluoroquinolones, cephalosporins and carbapenems. Patients infected with isolates resistant to all these antibiotics classes experienced a 40% increase in the adjusted mortality rates compared to those with susceptible strains.^29^ Similarly, MDR of *Acinetobacter* infections further complicates treatment. Patients frequently have prolonged exposure to healthcare environments, increasing risk of exposure to antibiotics and colonization by resistant isolates.^30^

### Study Goals and Outline

As stated above, WHO lists carbapenem-resistant *Acinetobacter baumannii* and *Pseudomonas aeruginosa* as critical and high priority pathogens, respectively, as they are common, fatal nosocomial infectious agents.^3,31^ This study aims to provide further evidence for the hypothesis that QSIs can synergize with existing antibiotic drugs, leading to novel combination therapies with the great potential to be effective against *A. baumannii* and/or *P. aeruginosa*. To achieve this goal, we focused on a knowledge-based discovery approach to identify QSIs reported in chemical bioactivity online databases and literature followed by Quantitative Structure Activity Relationship (QSAR) modeling to identify additional approved or investigational drugs predicted as QSIs. We endeavored to nominate and test the combinations of putative QSIs with known antibiotics against various strains of *A. baumannii* and *P. aeruginosa* to identify synergistic treatments.

## Materials and Methods

### ChEMBL Mining and Data Curation

We began the search for compounds that may inhibit QS by searching the ChEMBL database^32^ for assays containing the phrase “quorum sensing”. Then, assays specific to the “Pseudomonas aeruginosa” and “Acinetobacter baumannii” species were selected for further curation. Data collected from ChEMBL were curated using KNIME (v4.5.2). These entries were separated by the assay “Standard Type” field annotated by ChEMBL. We discovered 1007 “Inhibition”, 141 “Activity”, 78 “Efficacy”, 71 “IC50”, 38 “Ratio”, 8 “CFU”, 4 “FC”, 3 “EC50”, 3 “Ratio IC50”, and 1 “IC85” entry Standard Types. The “Assay Description” field for each entry was reviewed to ensure that each assay was measuring the *inhibition* or *antagonism*, rather than *induction* or *agonism* of QS activity. Next, potential QS inhibitors were selected by choosing only entries with “Inhibition”, “Activity”, or “Efficacy” Standard Values >= 50%, or those with an “Active” or “Dose-dependent effect” annotation in the “Comments” field. The Assay Descriptions for the included entries were again manually reviewed and only entries that describe assays using compound concentrations of 10 µM or less were kept. “IC50” and “IC85” Standard Type entries were included only if their Standard Values were <= 10 µM. The results of this data curation exercise included 49 entries describing 36 unique compounds with >= 50% inhibition of QS activity at concentrations of 10 µM or less. 838 entries describing 349 unique compounds were regarded as inactive with these criteria. Both chemical structures and biological activities were curated following the protocols developed by us earlier.^33,34^ The resulting dataset is included in **Supplementary Data 1**.

### QSAR Modeling and Virtual Screening

We employed QSAR modeling, validation, and virtual screening protocols implemented in KNIME (v4.5.2). The 36 active and 349 inactive compounds were aggregated as a training set for QSAR modeling. Compound structure cleaning and curation was performed. This included removal of salts, mixtures, and duplicates, as well as standardizing structure representations (aromatic rings, nitro groups, etc.).^33^ Following these procedures, 372 compounds (36 actives and 336 inactives) remained in the cleaned training set. We trained a random forest model and validated its performance using a five-fold external cross validation protocol. We then performed virtual screening of the Inxight pharmaceutical collection^35^ to identify approved or investigational compounds predicted to inhibit quorum sensing in *P. aeruginosa*. From the 36 known active QSIs and QSAR predicted actives, those which are already U.S. FDA approved drugs were selected for combination screening.

### Compound Selection Guided by Literature and Knowledge Graph Mining

Abstract Sifter (Version 5.5) is a publicly available Microsoft Excel workbook-based application that enhances PubMed’s search capabilities.^36^ The macro-enabled Microsoft Excel workbook, developed by the US EPA, was utilized to run a PubMed query to identify compounds that possess quorum-sensing inhibition against *A. baumannii*, since no relevant assay data was found in ChEMBL for this species. The keywords: “Acinetobacter baumannii” AND “quorum sensing” AND “inhibitors” were used. The search returned a total of 37 relevant articles. To further filter only the most relevant compounds demonstrating promising quorum sensing activity, all abstracts and papers were further investigated. If the compounds described in the research papers presented inhibition of >50% at 50 micromolar concentrations or less, the compounds were nominated for further testing, and the remaining compounds were removed.

In addition to Abstract Sifter, we used the ROBOKOP knowledge graph^37,38^ and ChemoText^39^ to find the additional evidence of nominated compounds being the QSI and/or their prior use as a constituent of antimicrobial mixture therapy. We also used these tools and QSAR models developed by us earlier^40^ to exclude the combinations that may lead to undesired drug-drug interactions or side effects.

### Bacterial Strains Tested

All strains of *P. aeruginosa*: MRSN 317 (NR-51516), MRSN 1344 (NR-51520), MRSN 1583 (NR-51524), PA14 (NR-50573), MRSN 315 (NR-51515) and *Acinetobacter Baumannii*: WC-136 (NR-19298), WC-487 (NR-19299), 137 (OIFC137) (NR-17777), BC-5 (NR-17783), and MRSN 1171 (NR-52153) were procured from ATCC.^41^ The selections of these resistant strains were based on their descriptions of the various resistant antibiotics.

### Antibiotic Testing of *P. aeruginosa* and *A. baumannii* in 1536-well Plate Format

LB Liquid cultures of both *P. aeruginosa* and *Acinetobacter Baumannii* strains were used for AC_50_ determination for individual antibiotics used in 1536 well plate format. Briefly, 4 µL of LB medium was dispensed into multi-well plates by a Multidrop Combi Reagent Dispenser (Thermo Scientific, Pittsburgh PA). Test compounds were delivered as a DMSO solution via a Kalypsys pintool transfer (San Diego, CA) and arrayed as eleven-point titrations, with final drug concentrations ranging from 46 µM to 0.18 µM. Following compound transfers, 2 µL of diluted overnight culture of *P. aeruginosa* were for overnight growth assay. The bacterial growth was assessed using a BacTiter-Glo Luminescent Cell Viability Assay (Promega, Madison, WI) by measuring the ATP quantity, which was directly proportional to the number of viable cells in the well. The luminescent signal was read with a ViewLux reader (Perkin Elmer, Norwalk, CT).

### Combinatorial Matrix Synergy Assay for *P. aeruginosa* and *A. baumannii* in 1536-well Plate Format

LB Liquid cultures of both *P. aeruginosa* and *Acinetobacter Baumannii* were utilized for predicted QSI and antibiotic mixtures in a matrix format in 1536-well plates. Briefly, 6 µL of LB medium was dispensed by a Multidrop Combi Reagent Dispenser (Thermo Scientific, Pittsburgh PA) into 1536-well solid bottom plates preplated with the selected compounds in combination using acoustic dispenser (Labcyte 650, Beckman Coulter, Indianapolis, IN) at six-point titrations. Final concentrations of the tested drugs ranged from 46 µM to 0.18 µM. Bacterial growth was monitored by assessing cell viability though OD at 600nm, which directly correlates with the number of viable cells in each well. Optical density readings were captured using Envision reader (Perkin Elmer, Norwalk, CT).

### Selection of Antibiotics Tested

We selected 21 antibiotics commonly used in clinics for treatments of bacterial infections based on a variety of mechanisms, targeting different aspects of bacterial growth and replication: Chlorhexidine, Ciprofloxacin hydrochloride, Cefepime, Enrofloxacin, Piperacillin, Tazobactam, Difloxacin, Grepafloxacin, Pazufloxacin (mesylate), Tobramycin, Meropenem, Ceftazidime, Aztreonam, Garenoxacin, Clinafloxacin, Trovafloxacin, Delafloxacin (meglumine), Avibactam sodium, Gemifloxacin, Epetraborole and Cefiderocol. Descriptions of common uses of these drugs, their mechanisms of action, and drug resistance potential are provided in **Supplemental Data 2**.

We screened our in-house antibiotic library including these 21 drugs and selected those showing activity against most strains for matrix screening/analysis (**Supplemental Data 3**). This led to the selection of 11 antibiotics used in combination matrix screening: Avibactam sodium, Cefepime, Ceftazidime, Chlorhexidine, Ciprofloxacin Hydrochloride, Difloxacin, Meropenem, Piperacillin, Tazobactam, Tobramycin, and Trovafloxacin.

## Results

**Figure 2.**
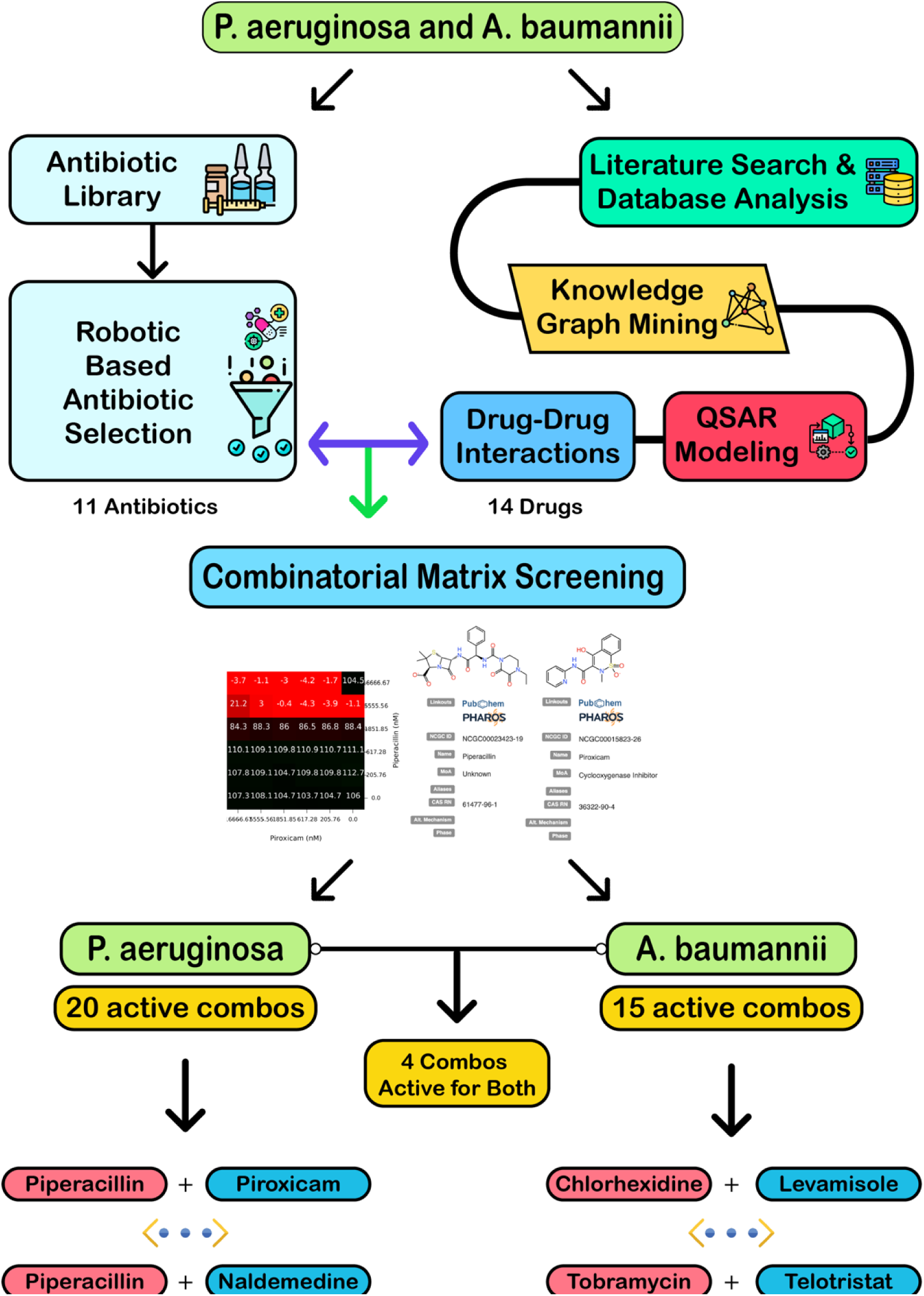
Study design and results of combinatorial matrix screening of antibiotics with QSIs. Antibiotics selected by robotic screening against drug-resistant bacteria and potential QSIs selected by knowledge mining and machine learning predictions were tested against *P. aeruginosa* and *A. baumannii* isolates, revealing synergistically active drug combinations.

### ChEMBL Quorum Sensing Inhibition Data

The result of the ChEMBL search yielded 1354 bioactivity entries for *P. aeruginosa,* but none were found for *A. baumannii*. The results of this data curation included 49 entries describing 36 unique compounds with >= 50% inhibition of QS activity at concentrations of 10 µM or less (**Supplemental Data 1**). 838 entries describing 349 unique compounds were regarded as inactive with these criteria. Of the 36 active compounds, 3 were chosen for synergy screening in this study due to their status as approved or investigational drugs in the Inxight pharmaceutical collection^35^: curcumin, azaguanine-8, and sulfathiazole.

### QSAR Modeling and Virtual Screening

The RF model trained on compounds tested for *P. aeruginosa* QSI activity obtained an AUC-ROC of 0.89 and the same model trained on randomly labelled data obtained performance of only 0.51, indicating our model was not overfitted.

**Figure 3.**
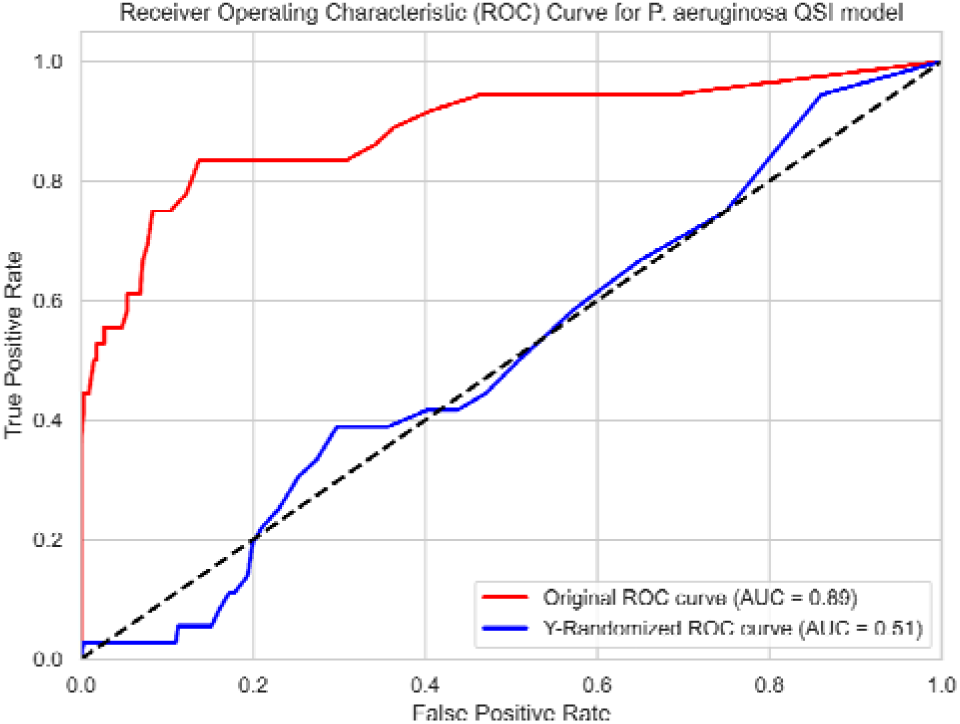
Receiver Operating Characteristic (ROC) curve of *P. aeruginosa* QSI classifier model demonstrating five-fold cross-validation performance. This model achieved 0.89 AUC (red line), while cross-validation on randomly labeled data achieved no better performance than random prediction (0.51 AUC, blue line).

We further performed virtual screening of the Inxight pharmaceutical collection.^35^ 49 of the 12,584 compounds were predicted to be active. Of these 49, seven compounds including abafungin, mycophenolic acid, telotristat, fenebrutinib, umbralisib (R enantiomer), relacorilant, and naldemedine, remained as these were predicted not to have undesired drug-drug interactions by our models and knowledge graph mining results. Due to a significant deficit of reliable experimental data, we also utilized KGs for searching for any additional evidence of QSI or antibacterial activity of selected compounds, similar to our previous antiviral studies.^42^

### Abstract Sifter for Literature Mining

Literature mining with Abstract Sifter resulted in 37 articles for investigation. From these 37 articles, 26 compounds of potential interest were identified (Compounds with inhibition greater than 50% for *A. baumannii* and clearly stated concentrations used). From these 26 compounds, known antibiotics were excluded, resulting in the nomination of 5 compounds for further investigation: ketoprofen, piroxicam, indomethacin, curcumin, and levamisole.

**Table 1:** Nominated Compounds from *A. baumannii* Literature Search. Ketoprofen, piroxicam, indomethacin, curcumin, and levamisole were identified as QSIs in *A. baumannii*.^43–45^

| Compound Name: | PUBCHEM ID | Inhibition | Concentration | DOI |
| --- | --- | --- | --- | --- |
| Ketoprofen | PUBCHEM.COMPOUND:3825 | 72% | 0.7-6.25 mg/mL | [10.1111/jam.15609](https://doi.org/10.1111/jam.15609) |
| Piroxicam | PUBCHEM.COMPOUND:54676228 | 91% | 1.25-2.5 mg/mL | [10.1111/jam.15609](https://doi.org/10.1111/jam.15609) |
| Indomethacin | PUBCHEM.COMPOUND:3715 | 81% | 3.12-12.5 mg/mL | [10.1111/jam.15609](https://doi.org/10.1111/jam.15609) |
| Curcumin | PUBCHEM.COMPOUND:969516 | 80% | 0.01 and 0.05 mg/mL | [10.3389/fmicb.2019.00990](https://doi.org/10.3389/fmicb.2019.00990) |
| Levamisole | PUBCHEM.COMPOUND:26879 | 72% | 0.512 mg/mL | [10.1007/s10096-020-03882-z](https://doi.org/10.1007/s10096-020-03882-z) |

### Synergy and Antagonism of Selected Combinations

Figure 4 summarizes the results of testing the combinations of Piperacillin with Piroxicam at different concentrations against *Pseudomonas aeruginosa* (MRSN 1344). The heat maps depict the response profiles when testing these combinations, indicating areas of synergy and antagonism. The response profiles in the heat maps are evaluated using the DBSumNeg value (Figure 4 (**b**)), which quantifies the degree of synergy. In this context, DBSumNeg represents the sum of the differences between observed and expected growth inhibition values, with negative values indicating synergy. A DBSumNeg threshold of −3 is considered indicative of synergy because it reflects a significant deviation from additive effects, meaning the combined effect of the drugs is greater than the sum of their individual effects.^46^

**Figure 4.**
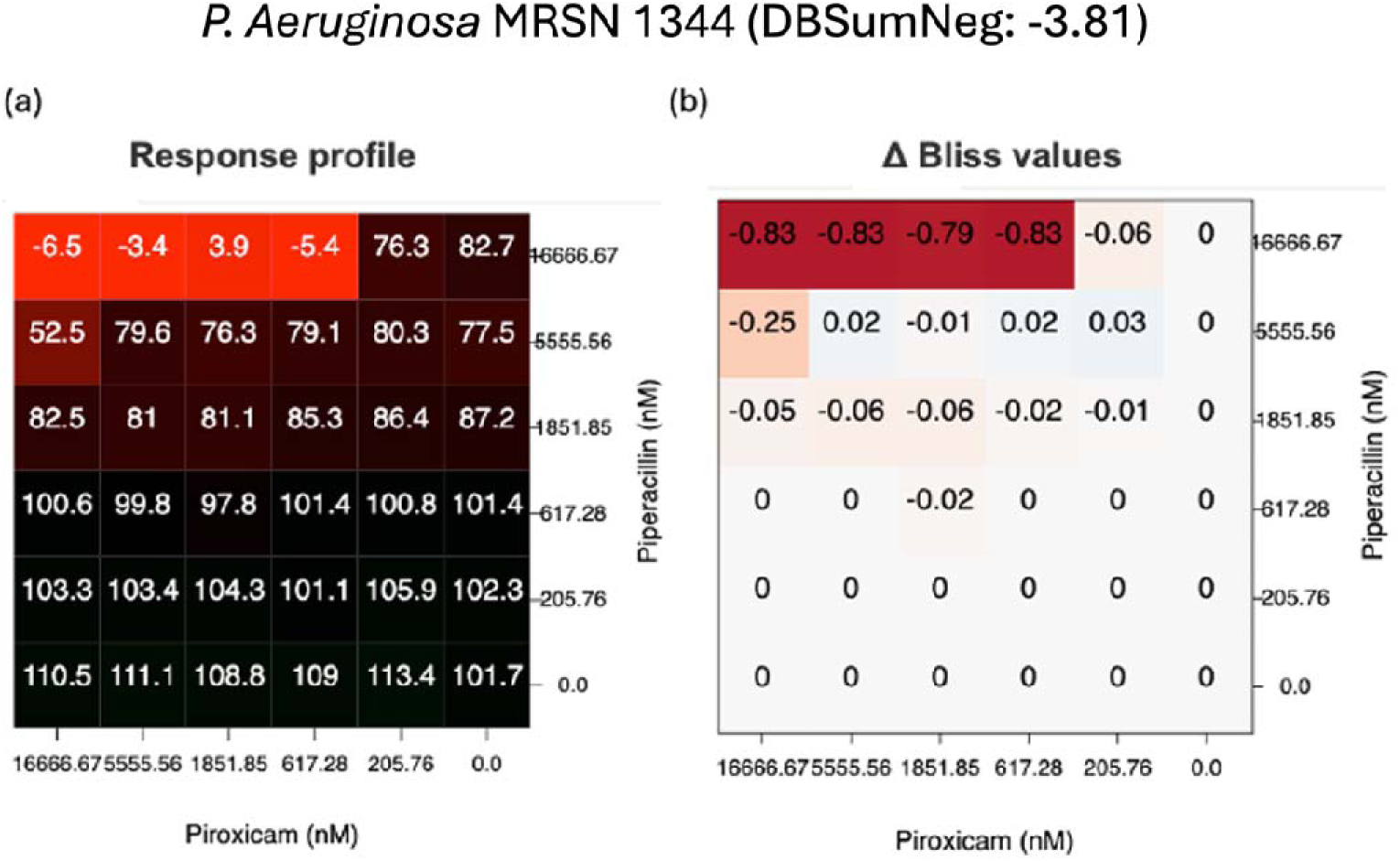
Representative Dose-Response Profiles for Piperacillin + Piroxicam against *P. aeruginosa* (MRSN 1344) DBSumNeg values are calculated from the combination of response profiles. Lower DBSumNeg Values (highlighted in red) represent stronger synergy.

Reviewing all response profiles for the drug combinations (**Supplemental Data 4-5**) resulted in selecting 26 combinations with DBSumNeg less than −3 in any tested strain (**Table 2**). Of these combinations, we sought to choose the combinations, for each bacterial species, which achieved the highest across the two tested species. We found that 22 combinations achieved synergy in one strain per species. The combination of <u>piroxicam and piperacillin</u> had strong synergy in both *A. baumannii* WC-136 and *P. aeruginosa* MRSN 1344, and the combination of <u>indomethacin</u> <u>and piperacillin</u> had strong synergy in both *A. baumannii* WC-136 and *P. aeruginosa* MRSN 1344. Combining <u>ketoprofen and piperacillin</u> had strong synergy in both *A. baumannii* WC-136 and *P. aeruginosa* MRSN 1344 (Figure 5).

**Figure 5:**
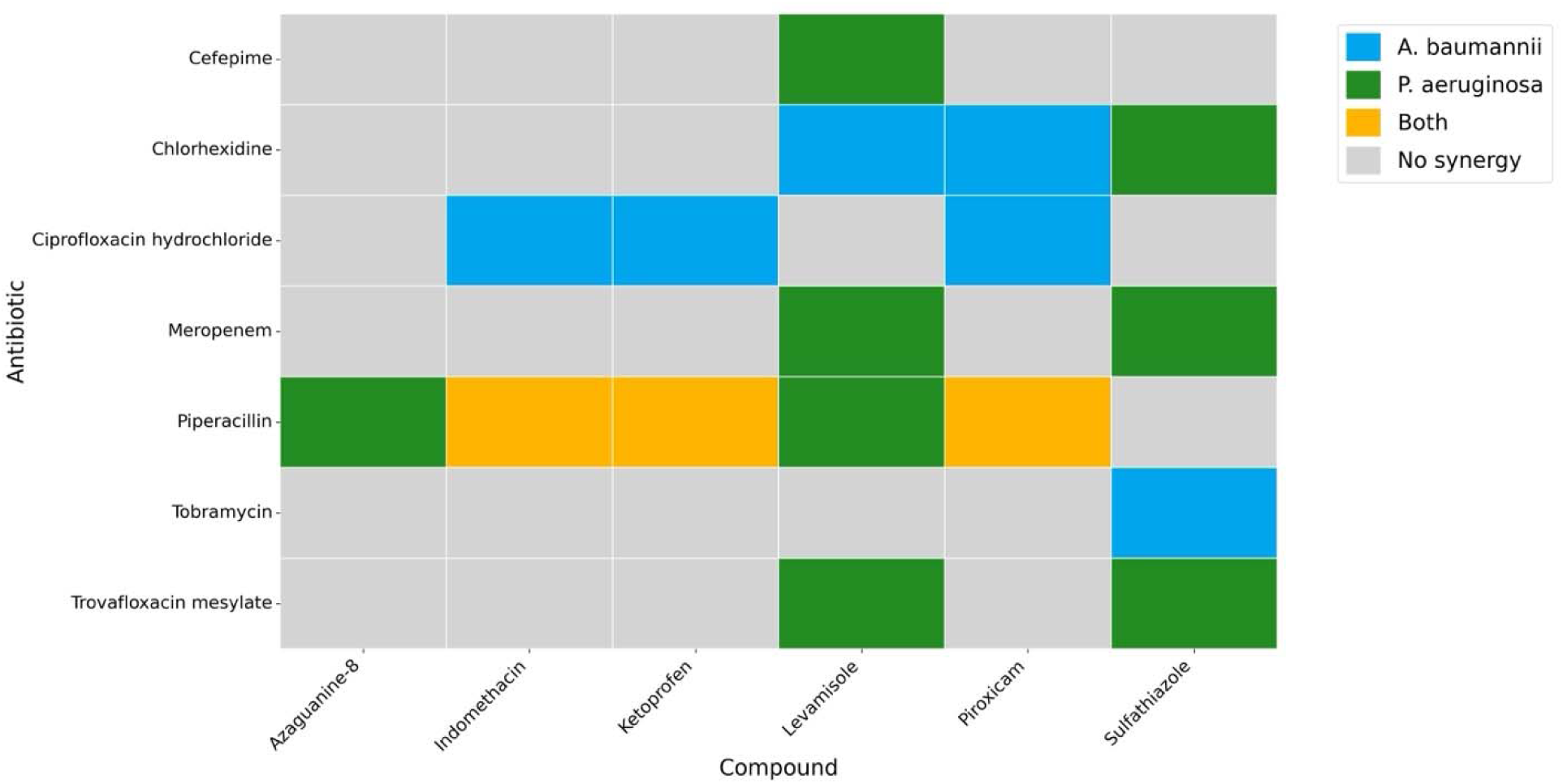
Synergistic Interaction Heatmap for Antibiotics and Compounds Against *A. baumannii* and *P. aeruginosa*. The figure displays a heatmap representing the number of strains showing synergy between various antibiotics and compounds, as evaluated against A. baumannii and P. aeruginosa. Number of strains showing synergy was quantified for combinations that showed a DBSumNeg value of less than –3.

**Table 2:**
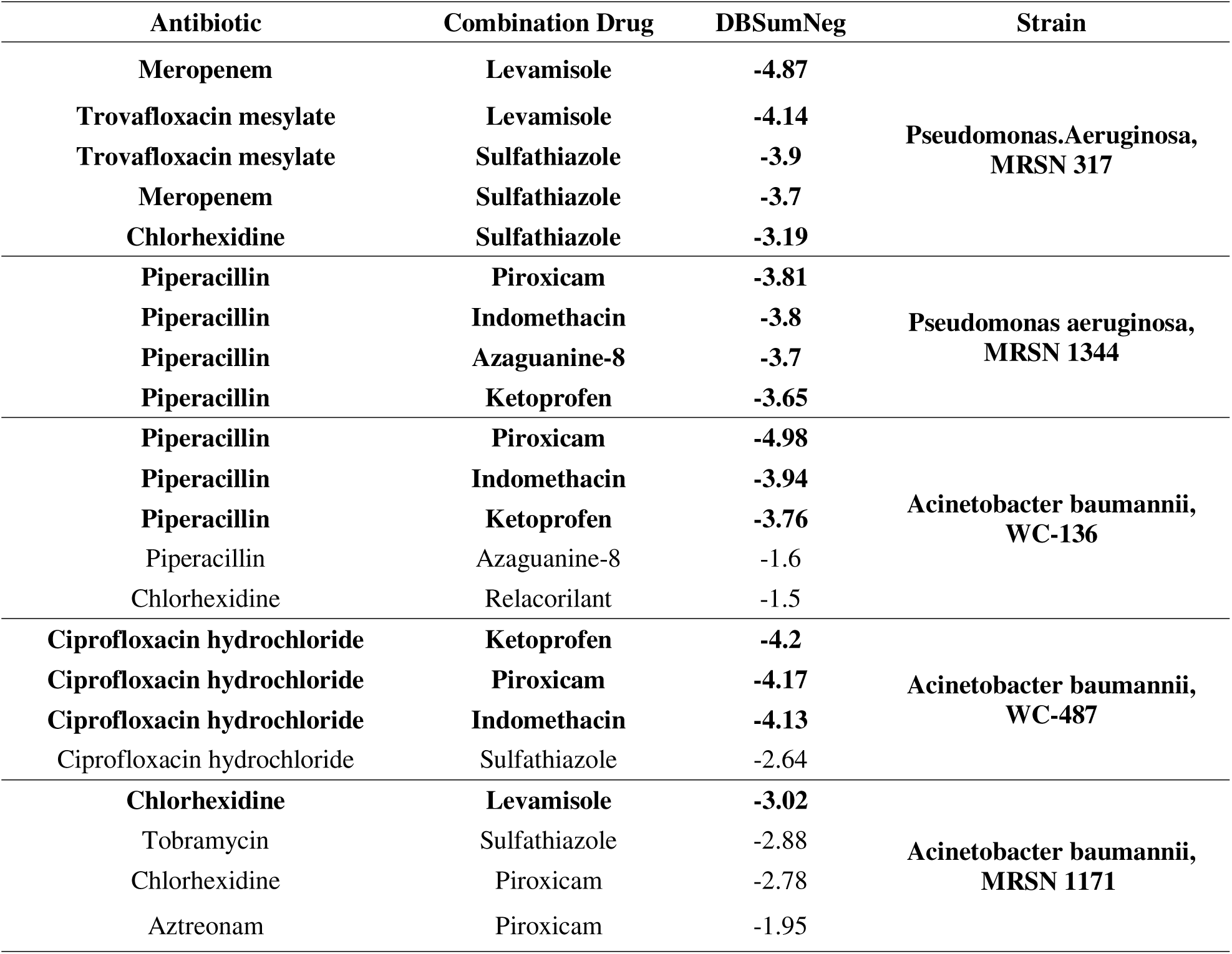
Combinations of known antibiotics with nominated drugs showing highest synergistic activity against *P. aeruginosa* and *A. baumannii* strains. Combinations with DBSumNeg less than −3.0 were selected for further review (drug names in bold font), but combinations with −2.5 or less are still shown.

We found that 27 drug combinations displayed anti-synergistic, antagonistic behavior in at least one strain. All but 4 of these antagonistic combinations were observed in only one bacterial strain. The combinations of azaguanine-8 and ciprofloxacin HCl, and abafungin and clinafloxacin showed antagonism in two *A. baumannii* strains. Figure 6 shows all synergistic and antagonistic combinations side-by-side in a circle plot for each bacterial species. A tabular form of these synergistic and antagonist interactions is available in **Supplemental Data 6.**

**Figure 6:**
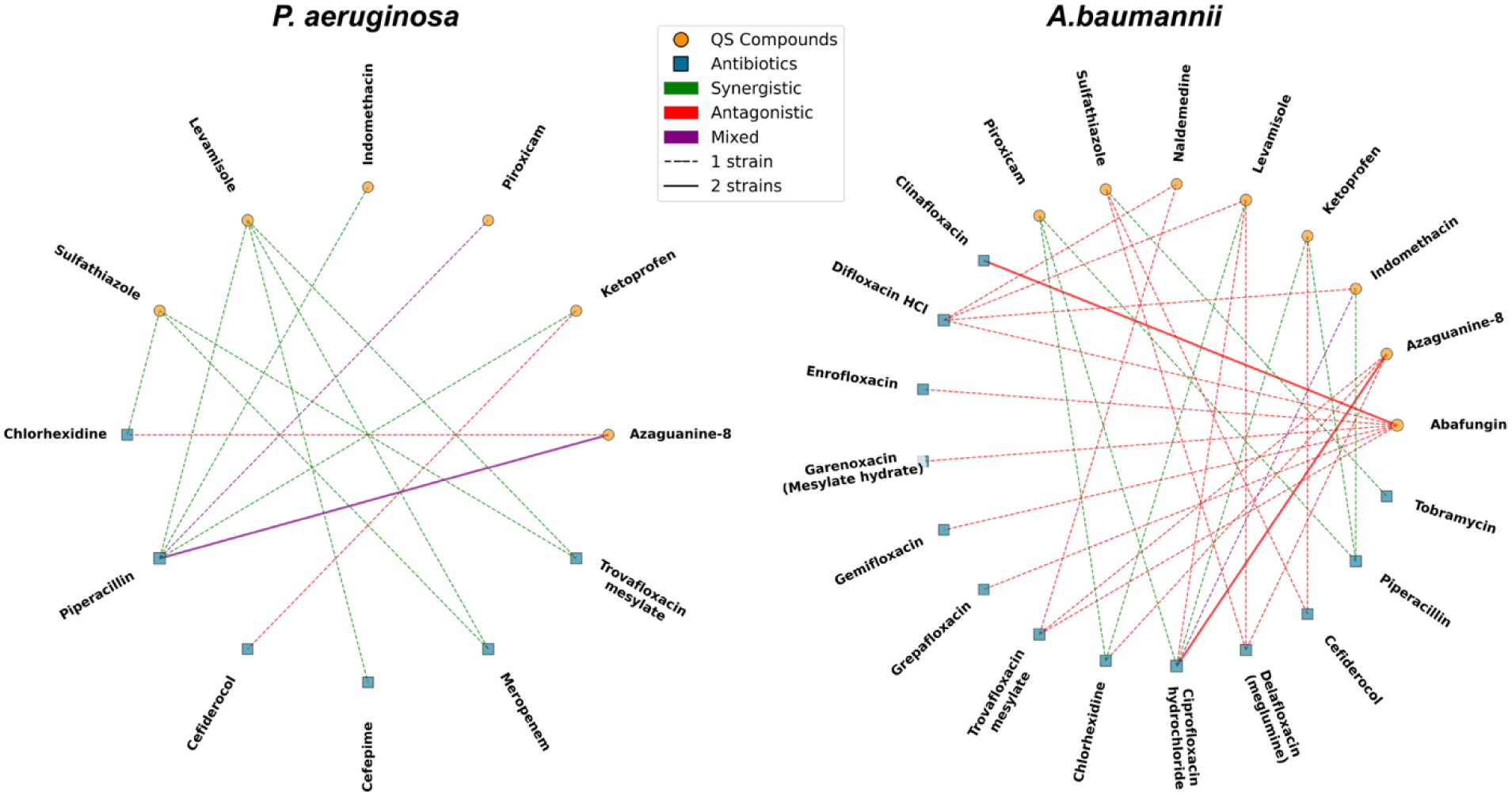
Interaction Network of *A. baumannii and P. aeruginosa* with Antibiotics and Compounds. Network represents the interactions between antibiotics and QSIs against *A. baumannii* (left) and *P. aeruginosa* (right), classified by their synergistic (green), and antagonistic (red).

### Drug Approval Status, Pharmacokinetics, and Dosing Considerations

We must also consider pharmacokinetic properties which might affect drug concentration at the site of infection. An important consideration is that the non-antibiotic quorum sensing inhibitors are not indicated for bacterial infections to date, so dosing regimens may need to be created to ensure the concentration reaches the IC50 in the site of infection, while mitigating potential toxicity. We have provided reported and predicted pharmacokinetic data for these drugs in **Supplemental Data 7.** We have not included CMax, as this is dependent on dosing and only directly applicable for bloodstream infections. Clinical use of these drug combinations is also contingent upon approval status of the drugs. Some of these drugs have either not yet been approved for clinical use in the US or may have been withdrawn due to safety concerns. Approval status of these drugs is compiled in **Table 3**.

**Table 3.**
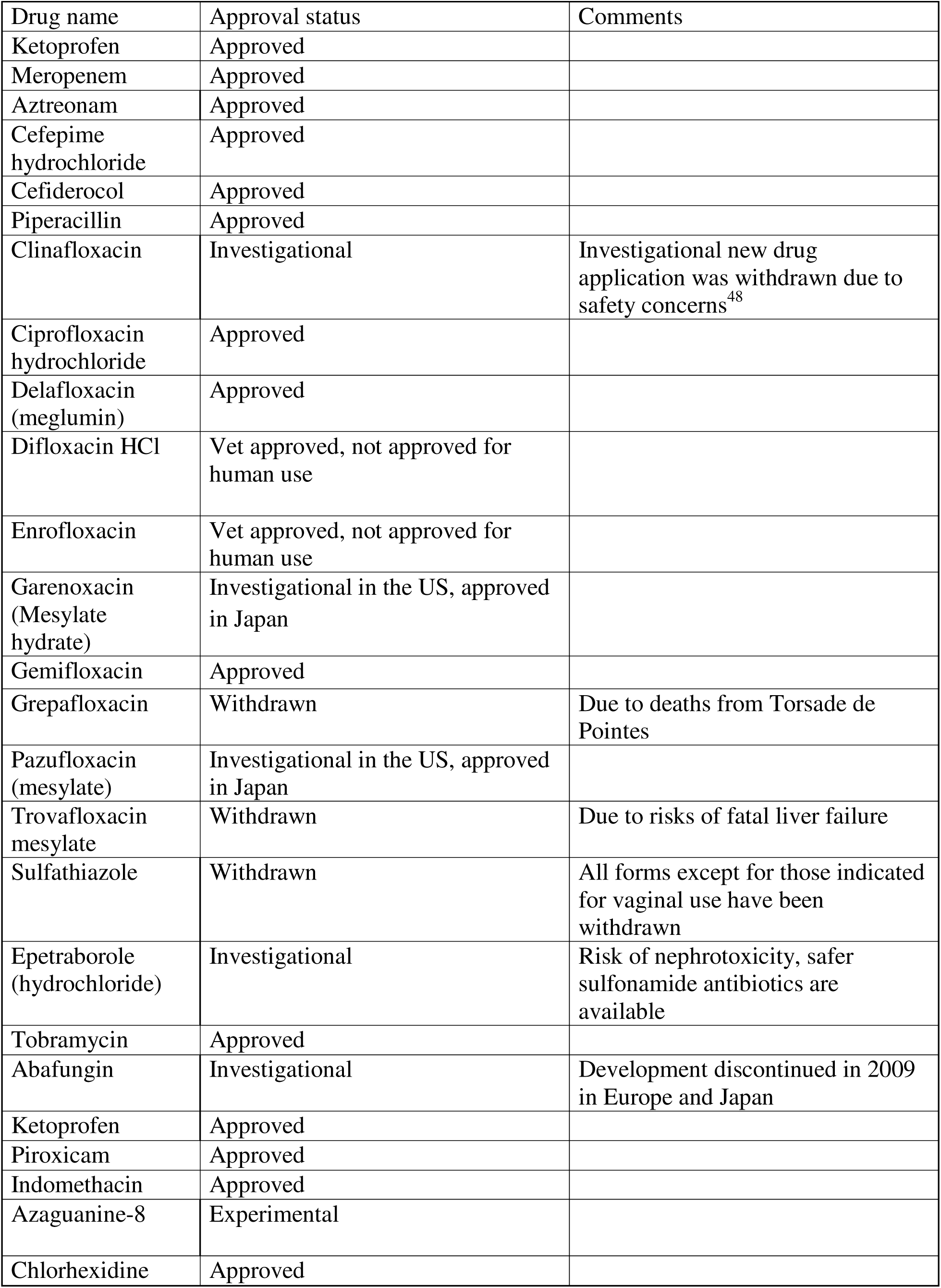
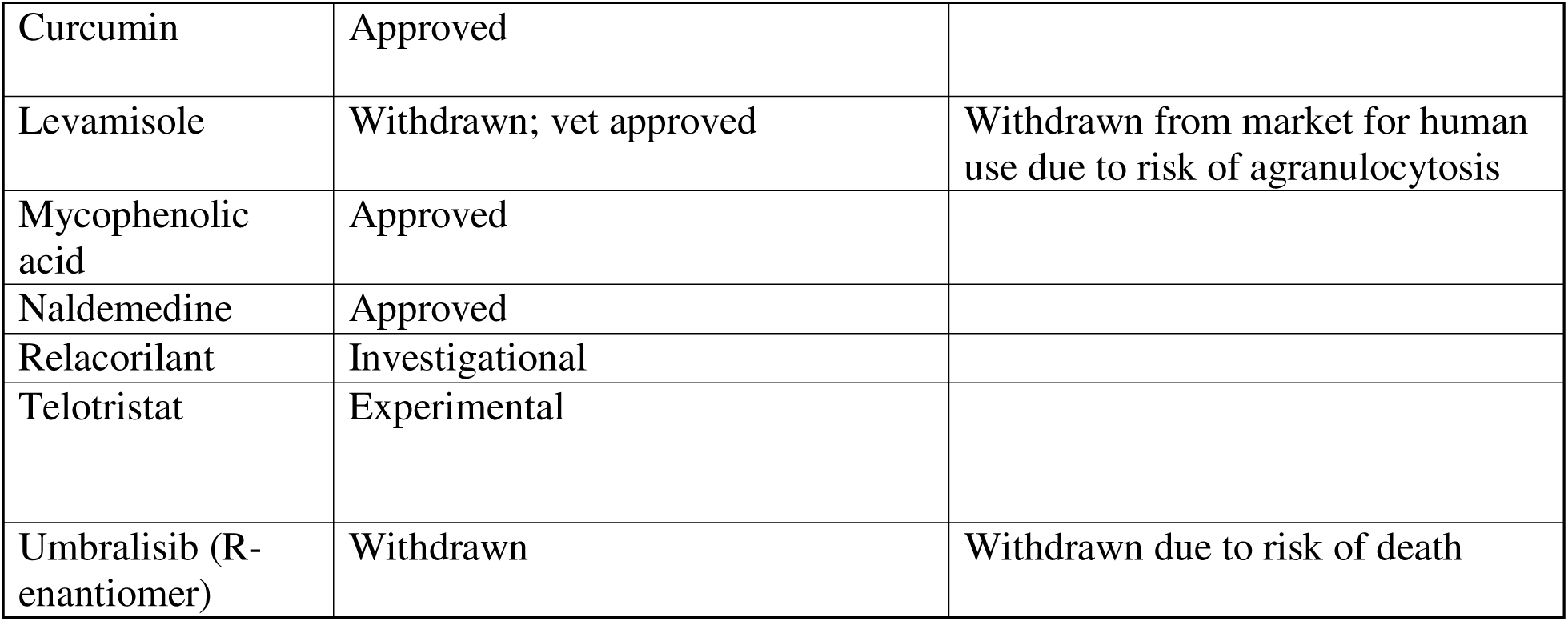
Approval status of tested drugs (found on DrugBank^47^)

**Tables 4a and 4b**. Potential interactions between selected synergistic combinations.

**Table 4.**
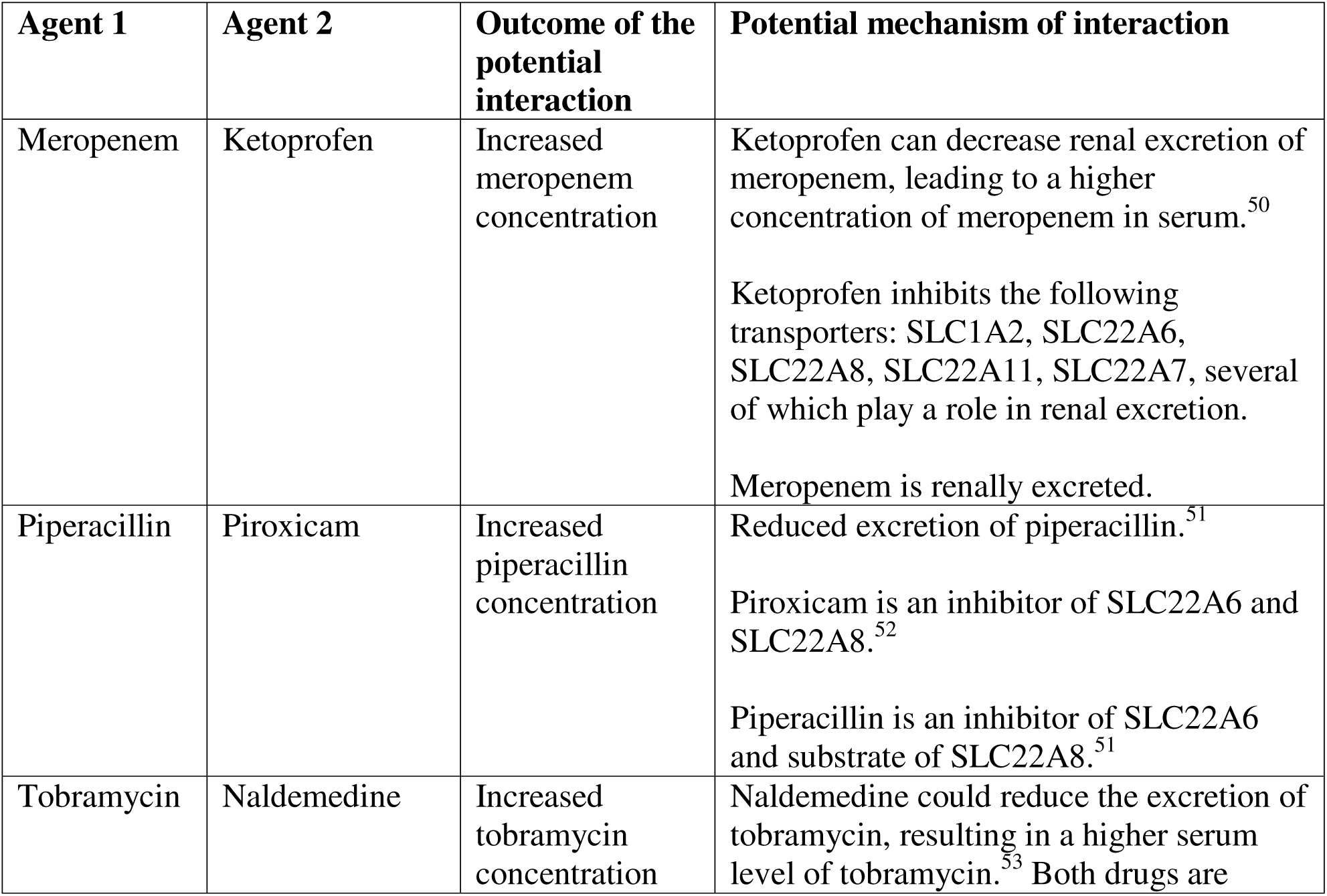
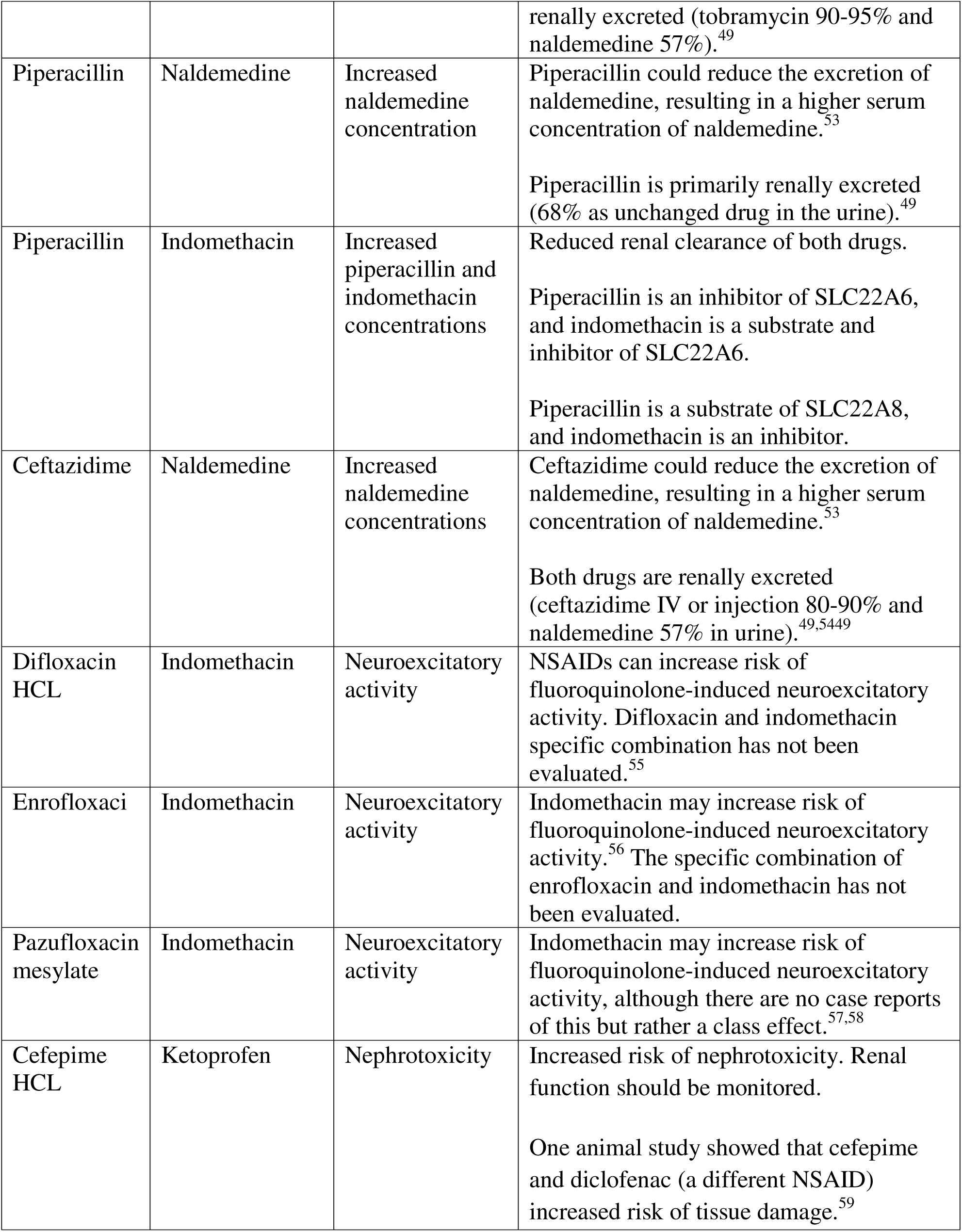
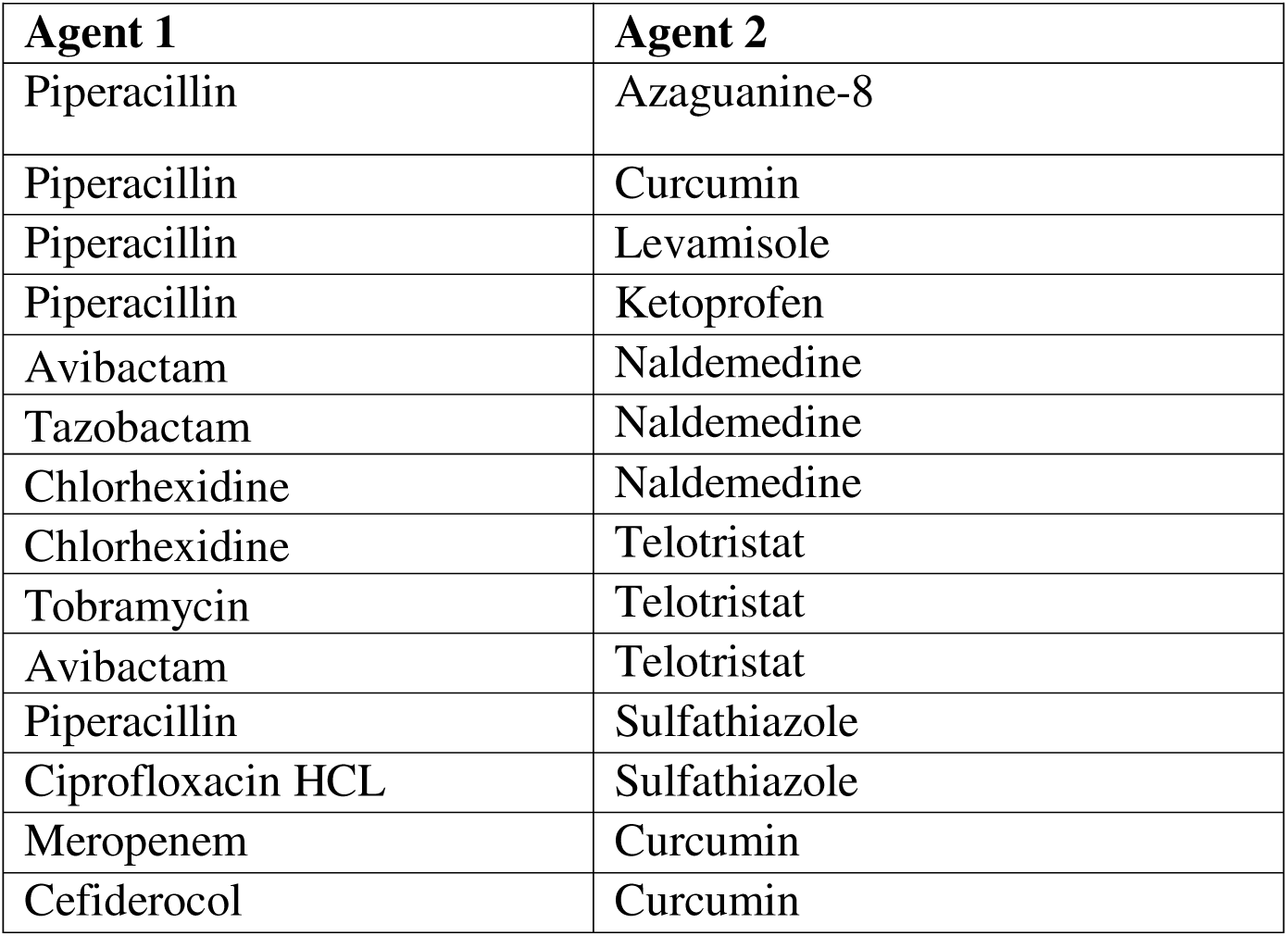
a. Drug combinations that are predicted to have drug-drug interactions. **b.** Drug combinations that have **no known** drug-drug interactions.

Interactions were checked using Facts and Comparisons^49^ and DrugBank^47^. If a potential interaction was found, we performed a further literature and database search for the nature of the interaction.

Of the 24 hit combinations, 10 are expected to have potential drug-drug interactions. Of note, many of these interactions pertain to one drug reducing the clearance of another, through inhibiting transporters (as in the case of ketoprofen + meropenem, piperacillin + piroxicam, and others). If these treatments are implemented in the hospital/clinic, dose adjustments may be necessary. However, there are several interactions more directly related to toxicity, as is the case with indomethacin + fluoroquinolones (risk of neuroexcitatory activity) and cefepime + ketoprofen (risk of acute kidney injury).

## Discussion

In this study, we substantiated the hypothesis that treating drug-resistant *P. aeruginosa* and *A. baumannii* isolates with QSIs in combination with existing antibiotic drugs could confer a synergistic bactericidal effect. To prioritize clinical translational potential, we chose to focus on screening combinations involving known QSIs which are also approved drugs with well-known safety and pharmacokinetic profiles in humans. To select additional test compounds with unknown QSI activity, we developed a QSAR model to predict QSI activity from among approved compounds in the Inxight pharmaceuticals collection. We identified several combinations involving piroxicam, indomethacin, ketoprofen, naldemedine, and telotristat which synergize with existing antibiotics. However, the exact mechanism of this synergistic activity has yet to be fully understood. Here, we discuss the putative mechanisms of action of these compounds based on prior literature.

### Putative Mechanisms of Action of the Synergistic QSIs

#### Piroxicam

While piroxicam has been shown to inhibit various QS virulence factors in several *A. baumannii* strains^43^, no studies, to the best of our knowledge, demonstrated its ability to inhibit QS, specifically, in *P. aeruginosa.* However, piroxicam was predicted to inhibit QS proteins LasR and PqsE by molecular docking and structure analysis.^60^ Piroxicam has been shown to protect mice from a lethal challenge of *P. aeruginosa* and diminish inflammatory response to *P. aeruginosa* pneumonia, although this protection was not due to direct bactericidal effects of the drug.^61^ Interestingly, other oxicam NSAID drugs, meloxicam and tenoxicam, have demonstrated QSI activity in *P. aeruginosa*.^62,63^

It is known that the combination of piroxicam with piperacillin can reduce the renal clearance of piperacillin. This is likely because piroxicam is an inhibitor of SLC22A6 and SLC22A8^52^, while piperacillin is an inhibitor of SLC22A6 and substrate of SLC22A8^51^. It is plausible that this drug interaction may improve treatment outcomes by extending the half-life of piperacillin, although additional *in vivo* studies are required to support this hypothesis.

#### Indomethacin

Like piroxicam, indomethacin has been shown to inhibit *A. baumannii* QS virulence factors, including biofilm formation, and surface motility, and bacterial tolerance to oxidative stress.^43^ In another study, indomethacin was tested for inhibition of QS in *Chromobacterium violaceum* CV026 and effects on virulence production in *Pseudomonas aeruginosa* PAO1, but was found to lack anti-QS activity as measured by the inhibition of production of violacein pigment.^64^ While we show the synergistic effects of indomethacin with piperacillin in our current study, these prior results suggest that indomethacin may be acting through mechanisms unrelated to QS. Further experiments beyond violacein pigment inhibition in *P. aeruginosa* are warranted to investigate the activity reported in our current study.

#### Ketoprofen

Ketoprofen, like piroxicam and indomethacin, was shown to inhibit QS virulence factors with MICs 0.7-6.25 mg/mL in various *A. baumannii* strains.^43^ The QSI activity of ketoprofen has also previously been confirmed in *P. aeruginosa* via assays showing attenuation of virulence factors and biofilm formation, as well as reduction in the expression of lasI, lasR, rhlI, and rhlR genes, by 35-47, 22-48, 34-67, and 43-56%, respectively.^65^ Furthermore, *in silico* studies and comparison of chemical structures to natural QS activator ligands suggest that ketoprofen and its analogues inhibit QS in *P. aeruginosa* by acting directly on the *PqsR* protein target.^66^

#### Azaguanine-8

Azaguanine-8 is a purine analog that has emerged as a potential quorum-sensing inhibitor in *P. aeruginosa*, identified through virtual screening for LasR.^67^ Tested in P. aeruginosa using a lasB GFP reporter assay, azaguanine-8 inhibited LasR controlled GFP expression in a dose-dependent manner with an IC of 0.64 µM, without affecting bacterial growth.^68^ The study by Tan et al. suggests that azaguanine-8 may have key hydrogen bond interactions with LasR residues Trp 60, Thr 75, and Tyr 93, consistent with a potential competitive antagonism mechanism, which may prevent receptor activation, leading to downregulation of LasR-controlled genes like B-elastase, and suppression of biofilm maturation and toxin secretion in *P. aeruginosa*.^68^

#### Sulfathiazole

Sulfathiazole is a sulfonamide antibiotic that acts as a competitive inhibitor of dihydropteroate synthase (DHPS) and an unconventional quorum-sensing inhibitor through indirect mechanisms. A high-throughput screen for anti-biofilm compounds in an *Escherichia coli* strain that overexpressed a diguanylate cyclase called AdrA identified sulfathiazole as a hit inhibiting DGC activity, lowering intracellular c-di-GMP levels, a well-known signaling molecule in bacteria that controls biological activity and biofilm production.^69^ *P. aeruginosa* is predicted to encode approximately 40 DGCs,^70^ with the only way to inactivate them all being to target shared motifs that generate c-di-GMB. Nonetheless, saturating the active site on various enzymes is impractical.

Alternatively, although direct binding of sulfathiazole to QS regulators (LasR, RhlR, PqsE/BfmR) has been scarcely observed, sulfathiazole-dependent changes to bacterial resistance caused by QS signal disruption may be connected to the modulation of genes in the folate biosynthetic pathway. Using quantitative reverse transcription PCR (qRT-PCR), folP expression was downregulated by 30% upon sulfathiazole challenge. Downregulation of folP, which codes for DHPS, may underlie the quorum-sensing inhibition potential of sulfathiazole.^71^

#### Levamisole

Levamisole is an anthelmintic drug repurposed as a potent QS inhibitor against both *P. aeruginosa* and *A. baumannii*. In vitro, *P. aeruginosa* treated at ⅛ of levamisole’s MIC attenuated various measures of QS-regulated phenotypes, decreasing static biofilm formation by 75%, swarming motility by 67%, and secretion of rhamnolipids by 53%.^45^ In a panel of multidrug-resistant *A. baumannii*, the same ⅛ MIC of levamisole achieved >50% inhibition of biofilm formation in 85% of strains. Further, using an *in vivo* mice mortality test, where they were injected with *A. baumannii*, treatment with levamisole improved survival from 0% to 60% at ⅛ MIC after 72 hours.

To understand the mechanism underlying levamisole action in *A. baumannii*, qRT-PCR revealed that levamisole (⅛ MIC) downregulated the AbaI gene, which codes for N-acyl-homoserine lactone (AHL)-synthase. Levamisole downregulated the AbaI gene by 83%, a key process for initiating the QS signaling pathway in *A. baumannii*. Without sufficient AHL, the AbaR receptor likely remains unliganded, and the QS pathway is silenced, blocking the coordinated expression of downstream virulence genes.

### Considerations for Clinical Use

Clinical use of these drug combinations will require consideration of drug distribution throughout the body and study of appropriate dosing strategies. Furthermore, clinicians must consider the half-lives of each combination therapy and devise dosing regimens to concomitantly ensure sufficient therapeutic concentrations of both drugs. Thus, additional monitoring may be required if these combinations are implemented. As usual, clinical judgment will be necessary for implementing these treatments in patients.

## Conclusions

In this study, we substantiated the hypothesis that treating drug-resistant *P. aeruginosa* and *A. baumannii* isolates with QSIs in combination with existing antibiotic drugs could confer a synergistic bactericidal effect. To prioritize clinical translational potential, we focused on screening combinations involving known QSIs, which are also approved drugs with well-known safety and pharmacokinetic profiles in humans. To select additional test compounds with unknown QSI activity, we developed a QSAR model to predict QSI activity from among approved compounds in the Inxight pharmaceuticals collection.

We used a combinatorial matrix screening approach to test these 14 identified compounds in combination with 11 antibiotics with antimicrobial activity against our *P. aeruginosa* and *A. baumannii* isolates at varied concentrations. We discovered several pairs of compounds with a potent antimicrobial synergy effect, including combinations of piroxicam, indomethacin, and ketoprofen with piperacillin, which showed synergy in both *P. aeruginosa* and *A. baumannii* strains.

We also investigated potential drug-drug interactions between the agents in the synergistic combination hits. Of the 24 combinations, 10 have potential drug-drug interactions. Of these, 6 pertain to the alteration in excretion. If implemented in the clinic/hospital, this could require dose adjustment, but it is not a reason to discount the combination. There are four combinations for which toxicity is a concern (NSAIDs + fluoroquinolones). These combinations could require further monitoring in the clinic.

In summary, our results and previous evidence support the hypothesis that QSIs could enhance the antimicrobial activity of the existing antibiotic arsenal. We anticipate these results will motivate further preclinical investigations to confirm antibacterial synergy and combination safety *in vivo* before translational use in the clinic.

## Conflict of Interest

A.T. and E.N.M. are co-founders of Predictive, LLC, which develops novel alternative methodologies and software for toxicity prediction. All the other authors declare no conflicts.

## Acknowledgements

This project has been funded in whole or in part with Federal funds from the National Institute of Allergy and Infectious Diseases, National Institutes of Health, Department of Health and Human Services, under Contract No. 75N93024C00038, awarded to Predictive LLC. AT and ENM also acknowledge the partial support from NIH (U24 ES035214) and NIEHS (R41ES033857).

## Notes

### Summary of Updates

Results and discussion and supplementary materials were revised to include the most recent experimental data.

