## Supplementary material for "Breaking the Phalanx: Overcoming Bacterial Drug Resistance with Quorum Sensing Inhibitors that Enhance Therapeutic Activity of Antibiotics": Supplemental Data 2 - Antibiotic Drugs Uses Mechanisms and Resistance.docx

1. Chlorhexidine

Usage: Antiseptic for skin disinfection before surgery and to sterilize surgical instruments. It is also used in mouthwashes to treat gingivitis. Mechanism of Action: Disrupts bacterial cell membranes and precipitates cell contents. Resistance: Rare, but overuse can lead to reduced effectiveness.

2. Ciprofloxacin Hydrochloride

Usage: Broad-spectrum antibiotic used to treat respiratory, urinary tract, gastrointestinal, and skin infections. Mechanism of Action: Inhibits bacterial DNA gyrase and topoisomerase IV, preventing DNA replication. Resistance: Increasing due to widespread use, especially in treating UTIs.

3. Cefepime

Usage: Broad-spectrum cephalosporin used for severe bacterial infections, including pneumonia, UTIs, and skin infections. Mechanism of Action: Inhibits bacterial cell wall synthesis by binding to penicillin-binding proteins (PBPs). Resistance: Emerging resistance, particularly in Gram-negative bacteria.

4. Enrofloxacin

Usage: Primarily used in veterinary medicine to treat bacterial infections in animals. Mechanism of Action: Inhibits bacterial DNA gyrase and topoisomerase IV, similar to ciprofloxacin. Resistance: Similar to other fluoroquinolones, resistance can develop with misuse.

5. Piperacillin

Usage: Antibiotic used for severe infections, including intra-abdominal and skin infections, and pneumonia. Mechanism of Action: Piperacillin inhibits bacterial cell wall synthesis; commonly paired with tazobactam, which inhibits beta-lactamases, extending piperacillin's spectrum. Resistance: Resistance observed in some Pseudomonas aeruginosa and Enterobacteriaceae strains.

6. Difloxacin

Usage: Fluoroquinolone used in veterinary medicine for bacterial infections. Mechanism of Action: Inhibits bacterial DNA gyrase and topoisomerase IV. Resistance: Similar concerns as other fluoroquinolones with potential resistance development.

7. Grepafloxacin

Usage: Previously used for respiratory and urinary tract infections; withdrawn due to safety concerns. Mechanism of Action: Inhibits bacterial DNA gyrase and topoisomerase IV. Resistance: Not applicable due to withdrawal.

8. Pazufloxacin (mesylate)

Usage: Broad-spectrum antibiotic used for bacterial infections, often in hospitalized patients. Mechanism of Action: Inhibits bacterial DNA gyrase and topoisomerase IV. Resistance: Emerging resistance patterns, particularly in hospital settings.

9. Tobramycin

Usage: Aminoglycoside used to treat severe bacterial infections, especially in cystic fibrosis patients. Mechanism of Action: Binds to the 30S ribosomal subunit, causing misreading of mRNA and inhibiting protein synthesis. Resistance: Can develop with prolonged use; often combined with other antibiotics to prevent resistance.

10. Meropenem

Usage: Broad-spectrum carbapenem used for severe and resistant bacterial infections. Mechanism of Action: Inhibits bacterial cell wall synthesis by binding to PBPs. Resistance: Carbapenem-resistant Enterobacteriaceae (CRE) are a significant concern.

11. Ceftazidime

Usage: Third-generation cephalosporin used for severe infections, including Pseudomonas aeruginosa infections. Mechanism of Action: Inhibits bacterial cell wall synthesis by binding to PBPs. Resistance: Resistance in Gram-negative bacteria, particularly Pseudomonas and Enterobacteriaceae.

12. Aztreonam

Usage: Monobactam antibiotic used for Gram-negative bacterial infections, especially in patients allergic to penicillins. Mechanism of Action: Inhibits bacterial cell wall synthesis by binding to PBPs. Resistance: Can occur, particularly in hospital-acquired infections.

13. Garenoxacin

Usage: Broad-spectrum quinolone used for respiratory and skin infections. Mechanism of Action: Inhibits bacterial DNA gyrase and topoisomerase IV. Resistance: Potential for resistance similar to other quinolones.

14. Clinafloxacin

Usage: Previously used for serious infections; withdrawn from the market. Mechanism of Action: Inhibits bacterial DNA gyrase and topoisomerase IV. Resistance: Not applicable due to withdrawal.

15. Trovafloxacin

Usage: Used for severe bacterial infections; withdrawn due to safety concerns. Mechanism of Action: Inhibits bacterial DNA gyrase and topoisomerase IV. Resistance: Not applicable due to withdrawal.

16. Delafloxacin (meglumine)

Usage: Fluoroquinolone used for acute bacterial skin and skin structure infections. Mechanism of Action: Inhibits bacterial DNA gyrase and topoisomerase IV. Resistance: Lower potential for resistance compared to older fluoroquinolones, but vigilance is necessary.

17. Avibactam Sodium

Usage: Beta-lactamase inhibitor used in combination with cephalosporins to treat resistant infections. Mechanism of Action: Inhibits beta-lactamase enzymes, preventing the degradation of beta-lactam antibiotics. Resistance: Resistance can develop, particularly with extended-spectrum beta-lactamases (ESBLs).

18. Gemifloxacin

Usage: Fluoroquinolone used for respiratory tract infections. Mechanism of Action: Inhibits bacterial DNA gyrase and topoisomerase IV. Resistance: Similar to other fluoroquinolones, resistance can develop with misuse.

19. Epetraborole

Usage: Investigational antibiotic with activity against resistant Gram-positive and Gram-negative bacteria. Mechanism of Action: Inhibits bacterial leucyl-tRNA synthetase, disrupting protein synthesis. Resistance: Still under study, but designed to target resistant bacteria.

20. Cefiderocol

Usage: Siderophore cephalosporin used for multidrug-resistant Gram-negative bacterial infections. Mechanism of Action: Inhibits bacterial cell wall synthesis by binding to PBPs and utilizes a siderophore mechanism to enter bacterial cells. Resistance: Effective against many resistant strains, but vigilance is necessary.

21. Tazobactam

Usage: Beta-lactamase inhibitor often combined with piperacillin to treat severe infections. Mechanism of Action: Inhibits beta-lactamase enzymes, preventing the degradation of beta-lactam antibiotics. Resistance: Resistance can develop in bacteria producing beta-lactamases that are not inhibited by tazobactam.
