## Supplementary material for "Breaking the Phalanx: Overcoming Bacterial Drug Resistance with Quorum Sensing Inhibitors that Enhance Therapeutic Activity of Antibiotics": Supplemental Data 7 - Pharmacokinetic Properties.docx

**Pharmacokinetic properties of top synergistic compounds.** Bioavailability (F), Biopharmaceutical Drug Disposition Classification System (BDDCS), Clearance (CL), Fraction excreted unchanged in urine (EoM), Fraction unbound in plasma (fu), Water solubility (S), Half-life (t_half), and Volume of distribution (Vd) data were collected from the DrugCentral database, unless otherwise cited.[^45^](https://sciwheel.com/work/citation?ids=3843257&pre=&suf=&sa=0&dbf=0)

| **Compound** | **F (%)** | **BDDCS** | **CL (mL/min/kg)** | **EoM (%)** | **fu** | **S (mg/mL)** | **t_half (hr)** | **Vd (L/kg)** |
| --- | --- | --- | --- | --- | --- | --- | --- | --- |
| Meropenem | 0 | 4 | 3.9 | 70 | 0.87 | 30[^47^](https://sciwheel.com/work/citation?ids=16453591&pre=&suf=&sa=0&dbf=0) | 1 | 0.3 |
| Piperacillin | 99 | 3 | 4 | 71 | 0.5 | 714.3 | 0.96 | 0.27 |
| Piroxicam | 99 | 2 | >1.00* | 0.5 | Binder* | 0.02 | 30-90 | 0.14 |
| Curcumin | Low[^56^](https://sciwheel.com/work/citation?ids=9394426&pre=&suf=&sa=0&dbf=0) | Not found | >1.00* | Trace amounts | Weak/nonbinder* | <0.01^ | 6-7 hr[^57^](https://sciwheel.com/work/citation?ids=2470678&pre=&suf=&sa=0&dbf=0) | Not found |
| Levamisole | 65 | 1 | >1.00* | Not found | 75-80 | 1.07^ | 4.4-5.6 | Not found |
| Indomethacin | 99 | 2 | 1.30 | 15 | 0.01% | 0.00 | 1.40 | 0.10 |
| Ketoprofen | 90 | 2 | 1.60 | 0.50% | 0.01% | 0.01 | 2.10 | 0.13 |
| Naldemedine | 20-56% | 1 | Not available and No prediction* | Not found | 0.06-0.07 | 1.14 | 11 | 155 L (total) |
| Telotristat | Above 80%* | Not found | 2.7 L/hr (telotristat ethyl); 152 L/hr (telotristat_ | Not found | >99% | 71 | 0.6 (telotristat ethyl); 5 (telotristat) | 428 L (total) |

*= predicted on PhaKinPro[^59^](https://sciwheel.com/work/citation?ids=16356908&pre=&suf=&sa=0&dbf=0)

^=predicted on StopLight[^60^](https://sciwheel.com/work/citation?ids=15311871&pre=&suf=&sa=0&dbf=0)
